# Motor planning under uncertainty

**DOI:** 10.1101/2021.02.11.430753

**Authors:** Laith Alhussein, Maurice A. Smith

**Affiliations:** John A. Paulson School of Engineering and Applied Sciences, Harvard University, Cambridge, Massachusetts, United States of America; Center for Brain Science, Harvard University, Cambridge, Massachusetts, United States of America

**Author notes:** Correspondence: (MAS).

## Abstract

Actions often require the selection of a specific goal amongst a range of possibilities, like when a softball player must precisely position her glove to field a fast-approaching ground ball. Previous studies have suggested that during goal uncertainty, the brain prepares for all potential goals in parallel and averages the corresponding motor plans to command an intermediate movement that is progressively refined as additional information becomes available. Although intermediate movements are widely observed, they could instead reflect a neural decision about the single best action choice at each point in time given the remaining uncertainty. Here we systematically dissociate these possibilities using novel experimental manipulations, and find that when confronted with uncertainty, humans generate a single motor plan that optimizes task performance, rather than averaging potential motor plans. In addition to accurate predictions of population-averaged changes in motor output, a novel computational model based on this performance-optimization theory accounted for a remarkable 80-90% of the variance for individual differences between participants. Our findings resolve a long-standing question about how the brain selects an action to execute during goal uncertainty, providing fundamental insight into motor planning in the nervous system.

## INTRODUCTION

We often plan actions before knowing the exact goal. For example, in baseball, a pitch can take as little as 400ms to reach the batter. With a reaction time of at least 200ms and a 150ms swing duration, the batter must initiate the swing based on visual information from only the first 10-20% of the ball’s flight, amidst considerable uncertainty surrounding the ultimate timing and positioning of the ball’s arrival at the plate. Nonetheless, batters deftly make use of this uncertain information and often make good contact with the ball. But what are the mechanisms that underlie this ability?

To study how motor planning proceeds when information is uncertain, controlled experiments have introduced uncertainty in a myriad of different ways: by delaying goal information^1,2^, pairing potential goals with distractor stimuli^3,4^, displaying noisy visual cues^5,6^, or incorporating high-level cognitive decision-making events during movement^7^. In response to this uncertainty, which prompts consideration of multiple potential goals, movements frequently begin in directions that are intermediate between them. These intermediate movements – widely considered a telltale sign of motor planning under uncertainty^1,5,8–14^ – are thought to provide fundamental insight into the neural processes by which the brain prepares an action to achieve a desired goal^1,3,8–10^.

The prevailing interpretation of intermediate movements during uncertainty is that they represent an average of the individual motor plans that arise from each potential target^1–3,8,11–27^. This idea, termed motor averaging, has been reported to occur for a number of key movement-related variables including a movement’s direction^1,2,20^, path shape^8,17^, effector orientation^18^, and even the feedback gains modulating how the motor system responds to errors induced by noise and external perturbations^15^. Moreover, motor averaging has been reported in a diverse array of behavioral paradigms, including point-to-point reaching arm movements^1,17^, saccadic eye movements^3,20,26^, and isometric force control^13,27^. In line with the motor averaging hypothesis, neurophysiological work has shown sensorimotor activity reflecting the parallel planning of movements to different potential targets^10,28–32^, with the assertion that parallel plans are automatically combined via motor averaging to form the final action plan^1,8,12,18,33^.

An alternative explanation for the occurrence of intermediate movements is that they instead reflect a single, deliberate plan that seeks to optimize task performance^5,34^. Unfortunately, a previous attempt to decouple this performance-optimization hypothesis from motor averaging was made only under conditions where intermediate movements between potential goals were rendered infeasible, using tasks with high-threshold movement speed criteria or with potential targets that had a wide spatial separation^9^. These tasks incentivize direct movements towards a single potential target for success, and thus promote a high-level explicit choice between targets before movement onset, rather than low-level motor planning given uncertainty. Such tasks are therefore unlikely to provide mechanistic insight into implicit motor planning.

It has been difficult to dissociate between motor averaging and performance-optimization because the motor plan that leads to task success often resembles an average of individual-target motor plans^5^. Indeed, close examination of previous studies that claim to provide support for motor averaging reveals not only that performance-optimization was not considered, but also that the observed results were, in fact, consistent with performance-optimization. For example, Chapman and colleagues (2010) examined motor planning during goal uncertainty by employing a task where participants were asked to make rapid reaching movements towards one of several potential target locations, with the final target cued only after movement onset (i.e., “go-before-you-know”). Analogously, Gallivan and colleagues (2016) studied motor planning at the level of feedback control policies in an analogous go-before-you-know task, but used targets of different widths in order to modulate the gain of feedback responses^35,36^. In both cases, intermediate actions were elicited when the goal was uncertain. In the Chapman study, the resulting movement was directed at the midpoint between the locations of the potential targets, and in the Gallivan study, the resulting feedback gain was sized at the midpoint between the gains associated with the potential targets. Although the authors interpreted these behaviors as evidence for motor averaging, in both cases the results can also be explained by performance-optimization: in the Chapman study, initial movements toward the spatial midpoint brings the hand closer to all potential targets, thus reducing the size of the subsequent movement required to reach the final target once identified, and in the Gallivan study, intermediate feedback gains balance the cost of the effort needed for movements executed with high-feedback gains, against the likelihood that this effort will be necessary.

In another example, Stewart and colleagues (2014) used the go-before-you-know paradigm to create goal uncertainty, but in combination with obstacles that were positioned to alter motor planning to one of the potential targets. In trials where the goal was initially uncertain, the obstacle placements resulted in intermediate movements that were deflected in a manner consistent with motor averaging, and this was taken as evidence for the motor averaging hypothesis. However, the obstacles were configured in such a way that the observed deflections on goal-uncertain trials, claimed to result from motor averaging, also happened to improve the safety margin around the obstacle. Therefore, performance-optimization for the goal-uncertain trials, which would result in motor planning that improved safety margins to minimize the likelihood of obstacle collisions, would also readily predict the experimentally observed deflections, calling the evidence for motor averaging into question.

The interpretational and methodological issues of the studies outlined above are emblematic of the pervasive confound between motor averaging and performance-optimization in studies examining motor planning under uncertainty. In the current study, we resolve the debate between these hypotheses by designing two different sets of experiments that allow rigorous dissociation of performance-optimization from motor averaging. In one, we employ obstacle-based perturbations of motor planning, like in Stewart et al. 2014, but using novel obstacles that break the congruence between the predictions for performance-optimization and motor averaging. In another, we create a novel dynamic environment that induces adaptive responses which make entirely opposite predictions for performance-optimization versus motor averaging. In both cases, we find clear evidence that, when faced with uncertainty, humans form a single motor plan that optimizes task performance.

## RESULTS

To decouple motor averaging (MA) from performance-optimization (PO) in both experiments (Expt 1 and Expt 2), we created modifications of the popular go-before-you-know task^5,8,9,12,15–18^, where uncertainty is introduced on some trials by presenting participants with multiple potential reach targets and disclosing the final goal location only after movement onset. We designed paradigms that altered motor plans in such a way that the MA and PO theories would make contrasting predictions when the reach goal was uncertain. In Expt 1, we accomplished this by pre-training a force-field (FF) environment that physically perturbed 1-target movements to left and right lateral target locations with one FF environment (FF_A_), but perturbed movements to a center target location with the opposite FF (FF_B_). Consequently, when the left and right target locations were presented as potential targets under uncertainty, MA predicts that intermediate movements incorporate the learned adaptive response to FF_A_. However, PO predicts that these intermediate movements should be planned so that they travel towards the midpoint of the potential targets in order to maximize the probability of final target acquisition. This would, in contrast to MA, predict that intermediate movements incorporate the learned adaptive response to FF_B_, appropriate for center-directed movements, allowing us to decisively dissociate PO from MA.

In Expt 2, we achieved a second dissociation between MA and PO by placing a virtual obstacle that induced deflections as it blocked direct movements to one target. Consequently, when that obstacle-obstructed target was used as a potential target under uncertainty, MA predicts that intermediate movements inherit a portion of this deflection, even if it steered them *closer to* the obstacle. In contrast, PO would predict that intermediate movements be consistently steered *away* from the obstacle to maintain a reasonable safety margin around it, as optimizing for task success would act to reduce the chance of obstacle collision. We thus present two distinct approaches that powerfully dissociate the predictions of the MA and PO theories for motor planning under uncertainty.

### Adaptation to novel physical dynamics can elucidate the mechanisms for motor planning under uncertainty

In Expt 1 (n=16), we employed a version of the ‘go-before-you-know’ task in which a combination of different FF environments was used to investigate movement planning under uncertainty. While gripping a robotic manipulandum that could apply forces to the hand (Fig 1a), participants initiated 20cm cued-onset reaching movements towards either a single pre-specified target (1-target trials; Fig 1b, *left*) or two potential targets (2-target trials; Fig 1b, *right*). On 1-target trials, the target, located at a left (+30°), right (−30°), or center (0°) eccentricity from the midline, was displayed for 1000ms before an auditory go cue was delivered. On 2-target trials, the same pair of potential targets always appeared in the left and right target locations for 1000 ms before the auditory go cue, but one (randomly selected) was extinguished 3cm after movement onset, leaving only the final reach target on-screen. Comparing the data from 1- and 2-target trials thus allowed us to infer how uncertainty about the final target location influences motor planning on 2-target trials.

**FIGURE 1.**
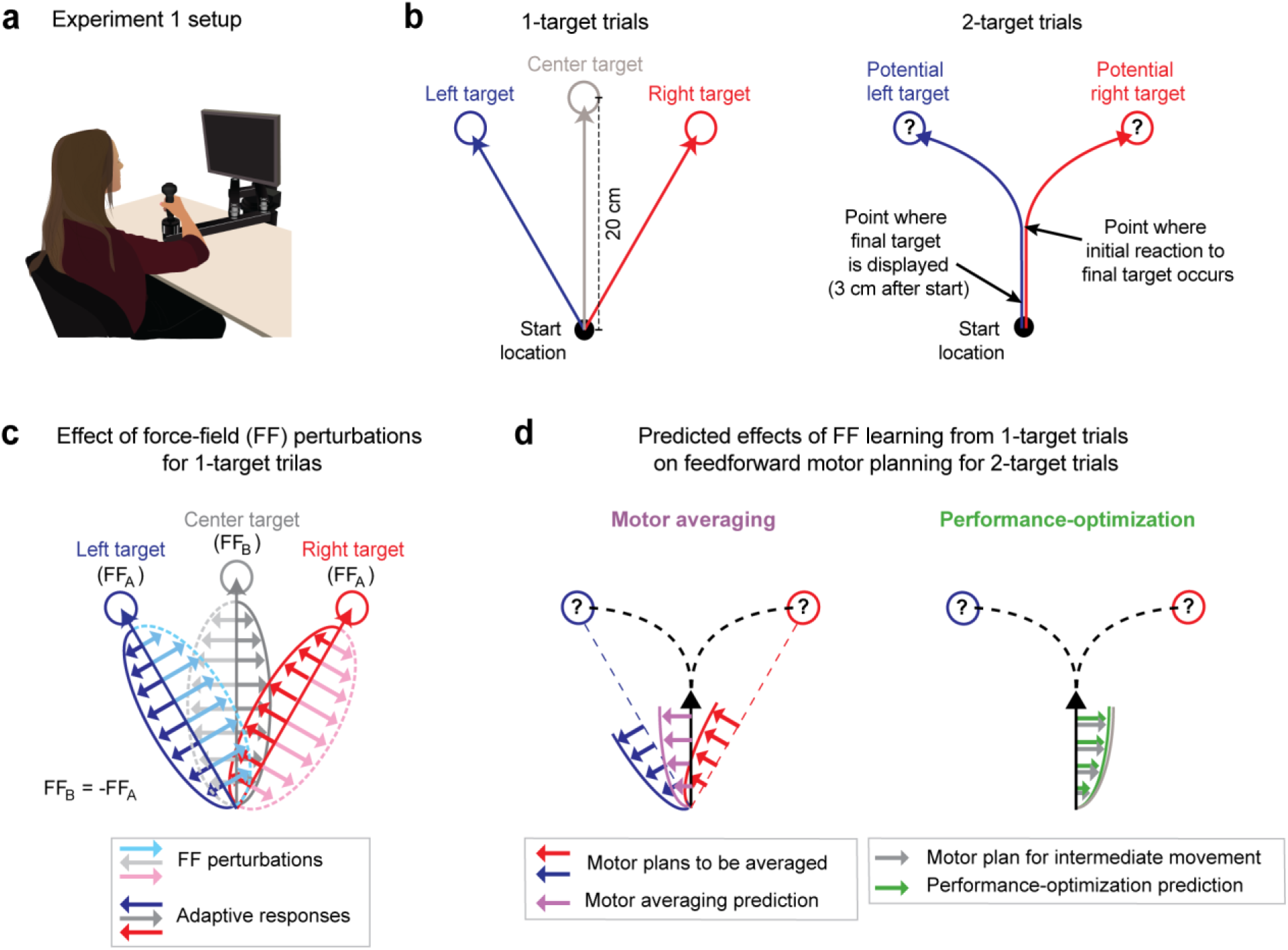
Multi-FF environment paradigm. **(a):** Setup for the multiple force-field (multi-FF) environment experiments. We altered environmental dynamics by exposing participants to viscous curl FFs in which the force vector perturbing reaching movements was proportional in magnitude and perpendicular in direction to the velocity of the hand (see methods). **(b):** Diagram of unperturbed 1- and 2-target trial types. On 1-target trials (*left* panel), a single target located in the left (+30°), right (−30°), or center (0°) direction was presented, and partcipants were instructed to initiate rapid 20cm reaching movements to the target after an auditory go cue. On 2-target trials (*right* panel), a pair of potenital targets, always in the left and right target directions, was presented before the go cue. Then, 3cm after movement onset, we extinguished one target and highlighted the other to indicate the final goal. Delaying the precise goal information in this manner typically leads to initial reaching movements directed in-between potential target locations before participants produce in-flight movement corrections toward the final target. **(c):** Individual target FF perturbations. During 1-target trials, we perturbed movements to the left and right targets with FF_A_ (light blue and pink arrow and dotted trace sets respectively) and the center target with FF_B_ (light grey arrow and dotted trace sets). The directions of FF_A_ and FF_B_ were counterbalanced across participants, but always with FF_B_=−FF_A_. Training in this composite environment alters the adaptive responses (darker solid arrow sets) in accordance with the FF imposed on each target. Note that the adaptive response on 1- and 2-target trials was measured as the force patterns participants produced on error clamp (EC) and partial error clamp (pEC) trials, respectively (see methods). **(d):** Predictions for motor averaging (MA) and performance-optimization (PO) for feedforward motor planning during uncertainty. Because both potential targets (left and right) were both associated with FF_A_, MA (purple arrows) predicts a force pattern consistent with FF_A_ on 2-target trials. However, since the initial motion on these 2-target trials is in the direction of the center target, PO (green arrows) predicts the force pattern consistent with FF_B_, which is opposite the MA prediction.

We designed a novel physical environment composed of multiple FFs that would result in very different predictions from MA vs. PO for motor planning on 2-target trials. Specifically, we trained participants to adapt to a viscous curl FF (see methods) that perturbed the left/center/right movements during 1-target trials with a FF_A_/FF_B_/FF_A_ composite environment (Fig 1c), where FF_B_ = −FF_A_, and the sign of the FFs was balanced across participants. For 2-target trials, MA would predict that participants average the force patterns (FF_A_ in both cases) learned for the left and right (lateral) targets, which correspond to the potential target locations on 2-target trials (Fig 1d, *left*). In contrast, PO would predict that participants produce the force pattern (FF_B_) appropriate for optimizing the planned intermediate movement since this movement maximizes the probability of successful target acquisition^5,34^ (Fig 1d, *right*). Importantly, and in contrast to previous studies^5, 12–14, 25–26^, we were able to reliably elicit intermediate movements during uncertainty in all our experiments by controlling the reward-rate with sliding scales for movement time criteria (see methods).

Participants first performed baseline 1- and 2-target trials to gain familiarity with them, and then completed a FF training block, where we imposed the multi-FF environment on 1-target trials as outlined above. Note that, because we differentially perturbed movements to the center target compared to the lateral targets, adaptation to the FFs associated with the lateral targets, FF_A_ in both bases, would interfere with adaptation to the FF associated with the center target, FF_B_. To elicit similar levels of FF adaptation for all three target locations, we included a greater number of training trials for the center target (using a ratio of 1:2:1 for left, center, and right target directions). The adaptive compensation to the FFs administered during training was measured with pseudorandomly interspersed error clamp (EC) trials^37^. After training, participants experienced a test epoch, which included 2-target partial error clamp (pEC) trials, on which the initial force pattern could be measured (see methods). The test epoch also included intermittent 1-target FF and EC trials, used to maintain the adaptive change in motor output developed during the training epoch and to measure its ongoing state.

### Adaptation to a multi-FF environment reveals that motor planning during uncertainty occurs via performance-optimization rather than motor averaging

Training within the multi-FF environment resulted in substantial levels of motor adaptation in all three target directions throughout both the training and test epochs as demonstrated in Fig 2a. We measured this adaptation with an adaptation coefficient (AC), a regression-based metric for comparing each movement’s force profile relative to the imposed FF perturbation (see methods)^38,39^. We found that the final AC levels achieved at the end of the training period (defined as the last 10% of EC trials) were within ~9% for all three target directions (0.73 ± 0.15 [95% C.I.], 0.80 ± 0.15, and 0.73 ± 0.12 for training in the left, center, and right target directions respectively). Importantly, these adaptation levels were largely maintained during the test epoch (0.71 ± 0.13, 0.68 ± 0.13, and 0.74 ± 0.09 for training in left, center, and right target directions respectively). Therefore, we used the population-averaged force profiles measured on 1-target EC trials (i.e., the adaptive response) during this epoch to construct our predictions for MA vs PO, as shown in Fig 2b. Note that all force data are aligned to target cue onset (T_ON_), the time at which one of the potential targets was extinguished for 2-target trials, which corresponds to the 3cm excursion point shown in Fig 1b. Also note that all force profile data are shown for the time that feedforward motor output was observed on 2-target trials, since forces due to feedback corrections do not reflect movement planning. We defined the time of feedback response onset (T_RESP_) as the point in time where we could detect statistically significant feedback responses during 2-target trials (150ms after target cue onset; see methods). In line with the AC measure we used to characterize overall learning, the more detailed analysis that we present of how force profiles evolve in time also normalizes raw force measurements by the level of the ideal compensation.

**FIGURE 2.**
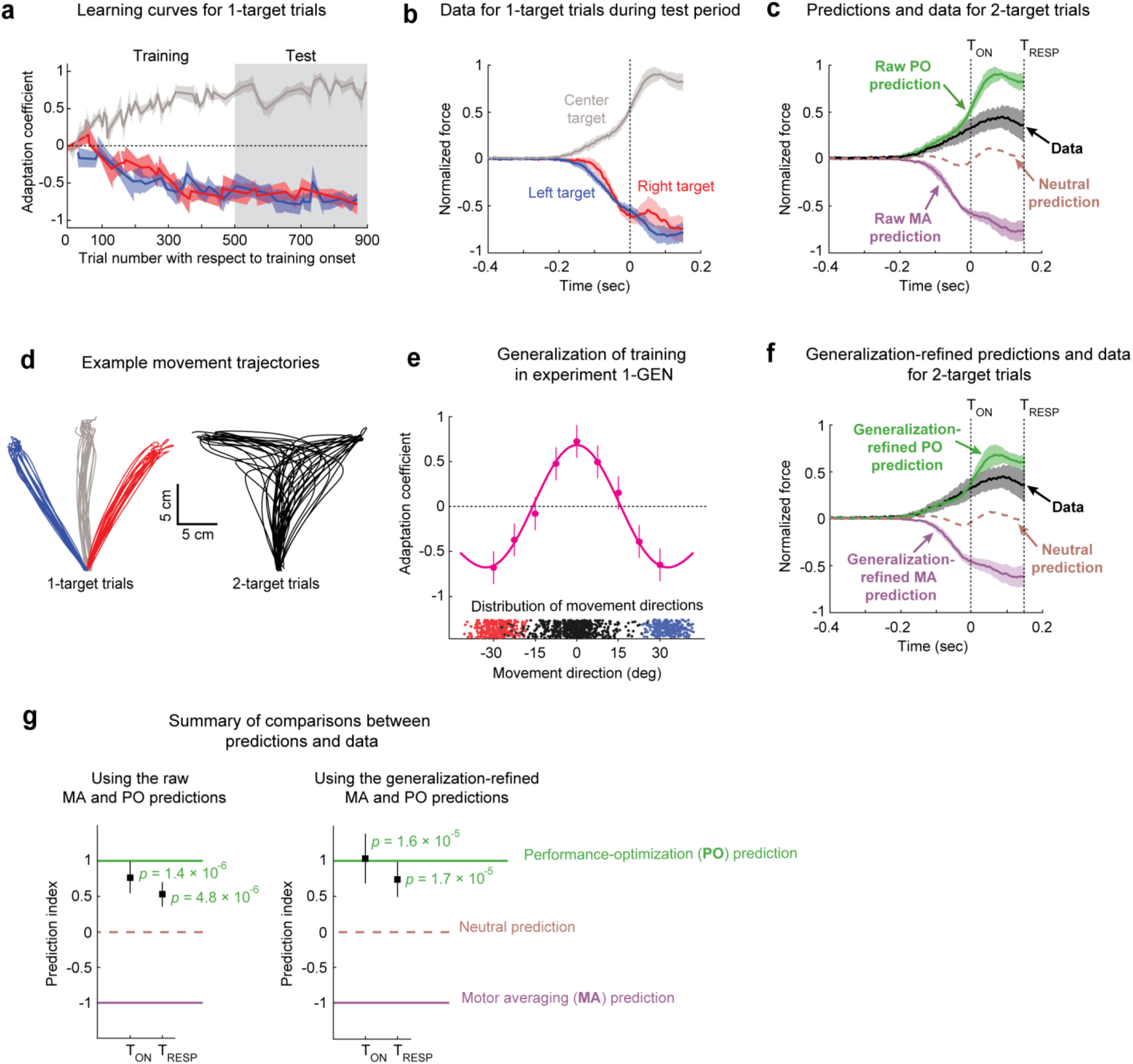
The effects of motor adaptation induced by novel environmental dynamics indicate performance-optimization during uncertainty. **(a):** Learning curves for 1-target trials. Participants (n=16) readily adapted left (blue), right (red), and center (grey) 1-target trial movements during the training period, and maintained this adaptation during the test period. Note that experiments were balanced with +15/−15/+15Ns/m FFs applied for the left/center/right targets in half the participants and −15/+15/−15Ns/m in the other half, and that adaptation levels of +1 refer to ideal learning for the center target FF, and adaptation levels of −1 refer to ideal learning for the left/right target FFs. **(b):** Population-averaged force profiles from 1-target EC trials measured during the test period. Data is normalized so that +1 refers to the amplitude of the maximum force in the ideal adaptive response for the FF associated with the center target. **(c):** Raw predictions (green & purple) and force profile data (black) for 2-target pEC trials. The raw PO prediction is based on the adaptive response for the center target, and the raw MA prediction is based on averaging the adaptive responses for the two (right and left) potential targets. The midpoint between the MA and PO predictions provides a neutral prediction reference (brown dashed trace), indicating that the experimental data are more closely aligned with PO than MA. **(d):** Randomly-selected trajectories from an example participant demonstrate non-trivial variability for movement directions in in both 1- and 2-target trials. **(e):** Generalization data (magenta) from Expt 1-GEN (n=10) is well characterized by a function obtained by summing gaussian generalization for each trained direction (+30°/0°/−30°). Blue and red dots indicate movement direction distributions for left and right 1-target trials and black dots indicate the corresponding distribution for 2-target trials across all participants. **(f):** Generalization-refined predictions and force profile data from 2-target pEC trials, where we used data from Expt 1-GEN to account for the effect of movement direction variability on the MA and PO predictions from Expt 1 (see methods). **(g):** Summary comparison between the data (black squares) and predictions (dashed lines). Both panels show a prediction index where −1 corresponds to the MA prediction and +1 corresponds to the PO prediction. This index was calculated by comparing the mean value for the force profile in the data to that from both model predictions, with means computed from the time of the go cue until either the target cue onset (T_ON_) or when target-specific responses were first statistically detectable (T_RESP_). The left panel shows results based on the raw predictions in panel **(c)** and the right panel shows results based on the refined predictions in panel **(f)**. The experimental data are significantly closer to the PO prediction than the nearly-opposite MA prediction, for both raw and generalization-refined versions of the predictions. In all panels, the shaded regions and error bars represent 95% CIs.

To specifically determine how movements are planned in uncertain conditions, we examined the force patterns predicted by MA and PO for 2-target trials, and compared them to the force patterns we measured on 2-target trials (Fig 2c). The MA and PO predictions are essentially opposite, consistent with the FF_A_/FF_B_/FF_A_ environment we imposed during training. The 2-target trial MA prediction corresponds to the average of the force profiles (adaptive responses) associated with the left and right 1-target EC trials plotted in Fig 2b, whereas the 2-target trial PO prediction corresponds to the force profile associated with the center target plotted in Fig 2b, as this is appropriate for optimizing a planned intermediate movement. Experimental data on 2-target trials show that participants systematically produce positive forces that are at odds with MA-based predictions, but in line with PO-based predictions for motor planning during uncertainty. We quantified the similarity between the 2-target trial data and the predictions using a prediction index (PI) that produces a value of +1 if the mean force from the experimental data were perfectly similar to the PO prediction, −1 if was perfectly similar to the MA prediction, and 0 if the data were halfway between both predictions (see methods). We measured this PI over two intervals: one extended for the duration we could reasonably use to examine feedforward motor output, spanning movement onset through feedback response onset (T_RESP_); the other was more conservative, spanning movement onset through target cue onset (T_ON_). In both cases, the PI estimates indicate that the observed 2-target trial force pattern is markedly more consistent with the PO than the MA prediction (T_ON_: PI = +0.77 ± 0.21 [95% C.I.], *p* = 1.41 × 10^−6^, t(15) = 7.25; T_RESP_: PI = +0.54 ± 0.16, *p* = 4.78 × 10^−6^, t(15) = 6.52; 1-sample *t-test*; Fig 2g, *left*).

### Refinement of model predictions based on movement direction variability

Observation of participant hand trajectories, exemplified in Fig 2d, reveals that there is appreciable variability in movement directions (*σ* ≈ 3.5° for 1-target FF trials and *σ* ≈ 5° for 2-target trials). This is important to consider because directional deviation from the intended target direction would lead to small but definite biases in the amount of adaptation for all three trained target locations (left/center/right), consequently biasing both the MA and PO predictions. Biases would occur, for example, because movements directed exactly in the 0° direction, towards the center target, would be associated with FF_B_ after training; however, off-target movements on either side of the 0° direction would be associated with a FF in-between FF_A_ and FF_B_. Thus, a distribution of movement directions centered around the 0° direction would, on-average, be associated with a FF in-between FF_A_ and FF_B_, rather than a FF comprised solely of FF_B_. This effect would be greater for movements farther off-target, resulting in a small but definite variability-dependent bias in the FF intended to be associated with a given target. We thus performed an additional experiment (Expt 1-GEN, n=10) where we measured the adaptation associated with a range of “off-target” movements to determine the size of this variability-induced bias (Fig 2e). Specifically, we employed a task design identical to Expt 1, with multi-FF training in the left/center/right target directions, but we replaced 2-target trials with 1-target EC trials and positioned them in directions that enabled a dense sampling for generalization of the trained adaptation (every 7.5° in-between the trained target directions). The resulting adaptation levels, shown in Fig 2e, show that the multi-FF environment generalizes nonlinearly across movement directions, with noticeable changes in adaptation around the trained target directions (a ~34% change 7.5° from the 0° trained target and 38-42% changes 7.5° from the ±30° trained targets).

The pattern of generalization observed in Fig 2e is well approximated (R^2^ = 0.98) by a model based on the additive combination of gaussians centered around the three trained target locations, in line with previous work examining the generalization of motor adaptation^40,41^. We therefore used this model to estimate the average adaptation level over the distribution of movement directions associated with left/right 1-target FF trials for the MA prediction and 2-target trials for the PO prediction from Expt 1. Each participant’s expected level of adaptation was then used to scale the raw force profiles that comprise the predictions. We found that this refinement reduced the magnitude of the predicted peak forces by 20-25% for both the MA and PO models (see Fig 2c vs 2f). Although the data were clearly more in line with the raw PO prediction than the raw MA prediction, there was still a noticeable mismatch between the PO prediction and the data (Fig 2c,g). However, taking generalization into account results in a refined PO prediction that is in even greater alignment with the data, corresponding to prediction indices that are closer to +1 at both T_ON_ and T_RESP_ (T_ON_: PI = +1.0 ± 0.4 [95% C.I.], *p* = 1.58 × 10^−5^, t(15) = 5.85; Fig 2g, *left*; T_RESP_: PI = +0.76 ±0.3, *p* = 1.72 × 10^−5^, t(15) = 5.80; 1-sample *t-test*; Fig 2g, *right*). Together these findings indicate that movements performed under uncertainty arise from a single action plan that optimizes task performance.

### Obstacle avoidance can elucidate the mechanisms for motor planning under uncertainty

In a second experiment, we used a different experimental approach to assess whether motor averaging (MA) or performance-optimization (PO) can explain the intermediate movements that arise from motor planning in uncertain conditions. We designed this experiment based on a subtle variation of an influential study that supported motor averaging^8^. However, here we show that the original instantiation of this experiment, which we replicate (Expt 2a), fails to make readily dissociable predictions for MA vs PO, but that a subtle modification of it (Expt 2b) leads to highly dissociable, and in fact essentially opposite predictions for these two models of motor planning under uncertainty.

In Expts 2a (n = 8) and 2b (n = 26), participants made 20cm cued-onset reaching arm movements (Fig 3a). We instructed participants to reach towards either a single pre-specified target or two potential targets, as in the 1- and 2-target trial designs from Expt 1 (obstacle-free trials; Fig 3b). After a baseline period in which participants practiced these trial types (see methods), we presented a virtual obstacle between the start position and, for example, the left target (Fig 3c-d). Please note that for simplicity, the subsequent explanations in this section reference a left-side obstacle condition; however, the experimental design was balanced within participants to include an equal number of interspersed right-side obstacle trials. In Expt 2a, we used an obstacle with the same size and positioning as that in Stewart et al. 2014 (ref. ^8^). This obstacle protruded 2cm to the right and effectively infinitely far to the left of the direct path between the start position and the left target (i.e., the obstacle-obstructed target), and thus required *rightward* deflections for left 1-target trials. In Expt 2b, we used a pared-down version of this obstacle that protruded 2cm to the right but 0cm to the left of the direct path between the start position and left target. This allowed for both leftward and rightward travel paths around the obstacle, but promoted less circuitous leftward deflections (see Fig 3c). Accordingly, all eight participants in Expt 2a consistently veered rightward as required around the obstacle, and all twenty-six participants in Expt 2b consistently veered leftward around the obstacle, in line with the intent of the experiment design. Hand paths from example participants are shown in Fig 4a, and data indicating deflections observed for all participants are shown in Fig 5b and 5e.

**FIGURE 3.**
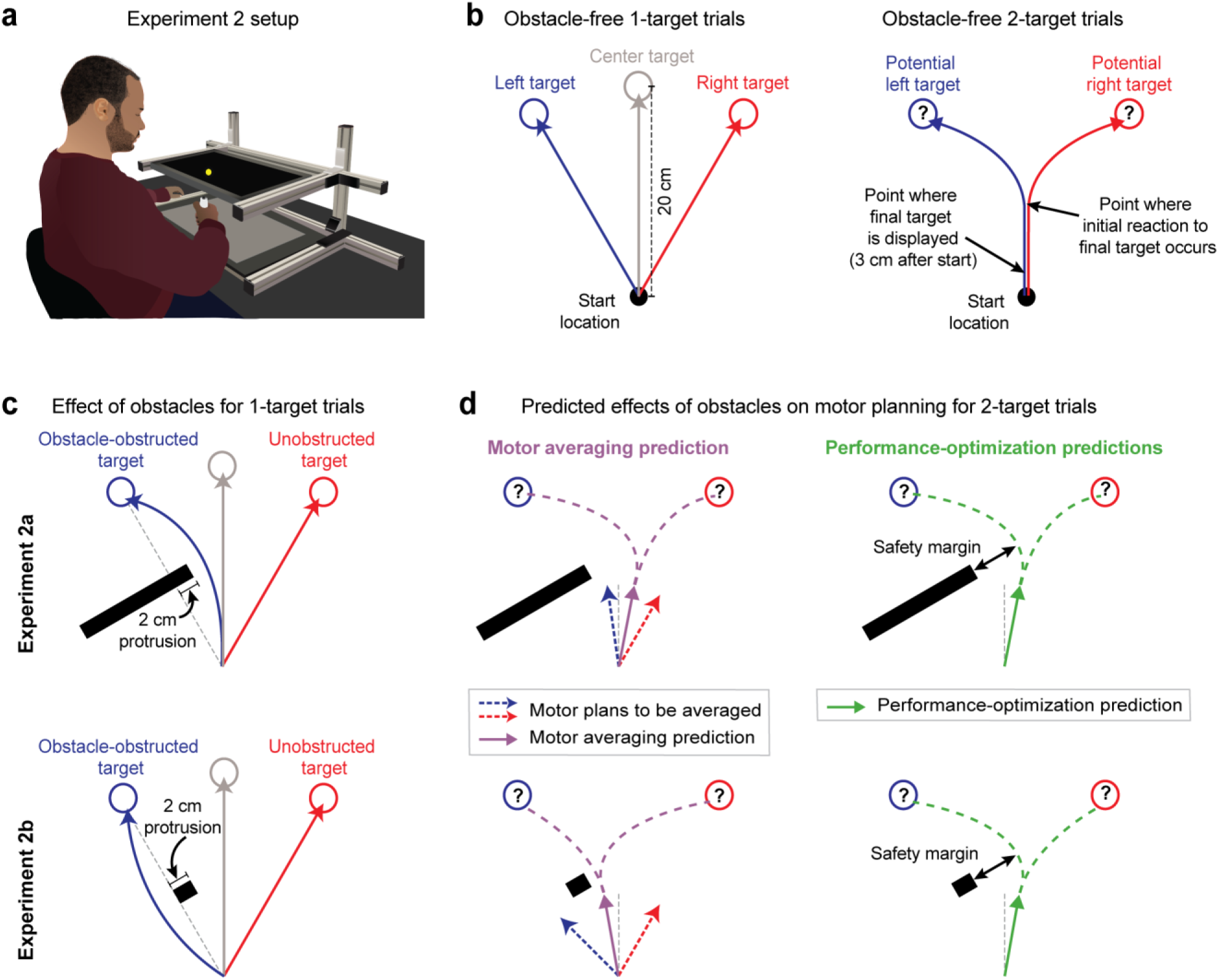
Obstacle avoidance paradigm. **(a):** Setup for the obstacle avoidance experiments, where virtual obstacles could obstruct movements to on-screen targets. **(b):** Illustration of obstacle-free 1-target and 2-target trials, similar to Fig 1b. **(c):** Diagram of left-side obstacle-present 1-target trials. The obstacle in Expt 2a was designed to elicit *rightward* movement deflections for left 1-target trials. The obstacle in Expt 2b, which included an identical protrusion towards the midline, was designed to promote a *leftward* deflection for left 1-target target trials. **(d):** Predictions during uncertainty. Left panels: MA (purple arrow) would average the plans for the obstacle-obstructed left and unobstructed right 1-target movements. It thus predicts initial movement directions (IMDs) on 2-target trials with opposite deflections from the midline for Expt 2a and Expt 2b (*away* from the obstacle in Expt 2a, but *towards* the obstacle in Expt 2b, where the safety margin around the obstacle would be reduced). Right panels: PO (green) would promote task success by balancing the costs of safely avoiding the obstacle and rapid target acquisition. It thus predicts IMDs on 2-target trials to be consistently deflected *away* from the obstacle. Thus, PO and MA make similar predictions for the obstacle in Expt 2a, but opposite predictions for the obstacle in Expt 2b. Note that **(c,d)** depict a left-side obstacle condition, but that the experiments were balanced within participants, with an equal number of trials for left and right side obstacles. A right-side obstacle condition would lead to mirror-opposite predictions for both MA and PO.

**FIGURE 4.**
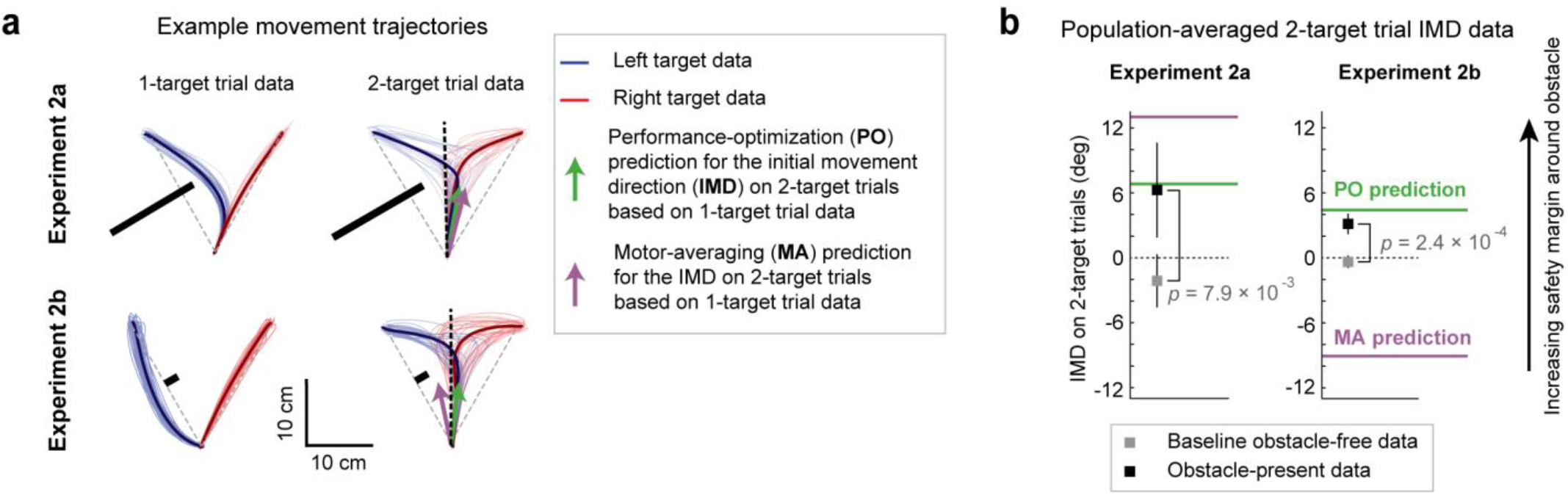
The effects of obstacle avoidance planning indicate performance-optimization during uncertainty. **(a):** Hand paths showing the effect of obstacles on 1- and 2-target trials for sample participants in Expts 2a-b. Thin lines indicate individual trial trajectories where the left (blue) and right (red) targets were cued, and bold traces indicate trial-averaged trajectories. On 1-targets, the obstacles induce substantial deflections for movements towards the obstacle-obstructed target, but in opposite directions for Expt 2a (n=8) and 2b (n=26). Motor averaging (MA) and performance-optimization (PO) predictions for the initial movement direction (IMD) on obstacle-present 2-target trials are displayed as purple and green arrows. These two predictions are similar in Expt 2a, but more distinct in Expt 2b. **(b):** Comparison of population-averaged IMDs on obstacle-free (grey squares) and obstacle-present (black squares) 2-target trials to model predictions by MA (purple line) and PO (green line). Results indicate that 2-target trial IMDs were significantly were biased away from the obstacle, closer to those predicted by PO rather than MA in both Expts 2a and 2b. Error bars represent 95% CIs.

**FIGURE 5.**
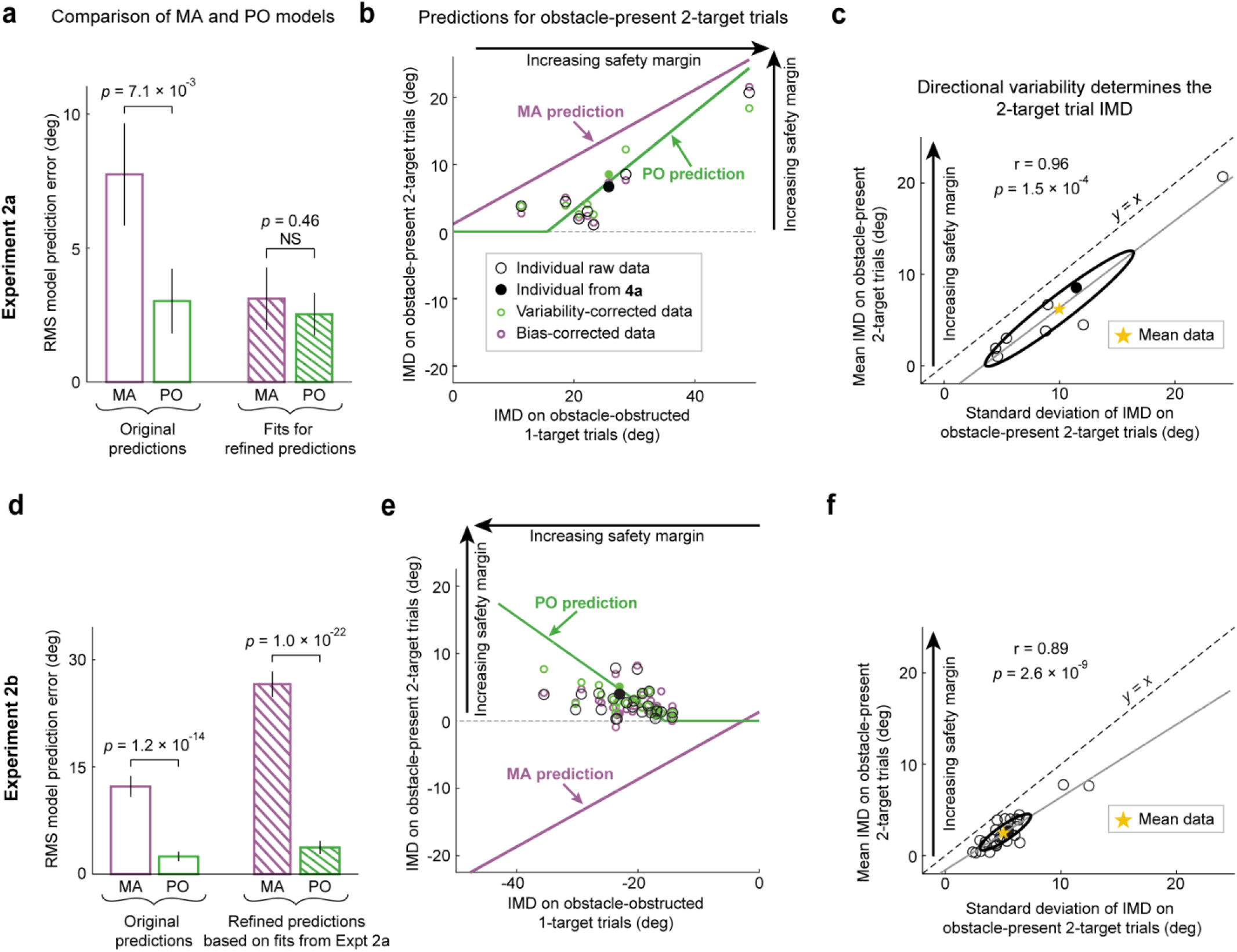
Performance-optimization predicts individual differences in obstacle avoidance. Analysis of initial movement directions (IMDs) on obstacle-obstructed 1-target trials and obstacle-present 2-target trials from Expts 2a-b. **(a-c)** show analogous analyses to **(d-f)**, for Expts 2a and 2b, respectively. **(a,d)** root mean square errors (RMSEs) for comparing model predictions to experimental data. For both models, we compared the original predictions (Eq. 1–2), which did not include any free parameters, and refined predictions, which included a single free parameter for each model. For MA, the original prediction assumes equal weighting between the obstacle-obstructed and unobstructed targets for motor averaging, whereas the refined prediction permits differential weighting. Analogously, for PO, the original prediction assumes equal weighting for the two determinants of task success: obstacle avoidance and movement timing, whereas the refined prediction permits differential weighting. For both the MA & PO refined models, we estimated the differential weighting using the Expt 2a data and tested it with the Expt 2b data. Thus the RMSEs based for the refined models in **(a)** are from *fits*, while those in **(d)** are from *predictions*. Results indicate that the original PO model predicts the data better than the original MA model in both Expts 2a and 2b, and that the refined PO model predicts the data better than the refined MA model in Expt 2b but parameter fits for these models in Expt 2a can explain the data equally well. **(b,e)** IMDs predicted for obstacle-present 2-target trials based on the IMDs observed on obstacle-obstructed 1-target trials, for PO (green trace) and MA (purple trace) hypotheses, compared to individual participant data. The MA and PO models predict similar IMDs in Expt 2a, but opposite IMDs in Expt 2b, allowing Expt 2b to provide a more powerful dissociation between them. Filled circles identify the participants shown in Fig 4a, and hollow circles show the remaining participants. Note that, although predictions are plotted with respect to IMDs observed on obstacle-obstructed 1-target trials, the MA model includes an additional dependency on IMDs observed for movements towards the unobstructed target and therefore, we corrected the raw data (black circles) to visualize this effect (purple circles). Analogously, the PO model includes a dependency on IMD variability, and we corrected the raw data for this effect as well (green circles). **(c,f)**: The relationship between the mean and standard deviation (SD) of IMDs on obstacle-present 2-target trials. Grey lines show linear fits to the data and confidence ellipses show 1 SD. A safety margin dependent PO hypothesis makes the key prediction that participants would size the mean IMD during uncertainty in proportion to directional variability to maintain a fixed statistical confidence against obstacle collision. In contrast, the MA hypothesis predicts no relationship between these variables. Both Expt 2a **(c)** and Expt 2b **(f)** show strong positive relationships between the mean and SD in the data (*r*=+0.96 and +0.89), consistent with the PO prediction but at odds with the MA prediction. Stars indicate mean across participants. Error bars represent 95% CIs.

The contrasting movement deflections produced by the different obstacles on 1-target trials in Expt 2a vs Expt 2b result in contrasting predictions for the MA hypothesis in these two experiments. On 2-target trials, where uncertainty in motor planning is present, MA predicts that participants would average the motor plans associated with obstacle-obstructed left 1-target trials and the unobstructed right 1-target trials,

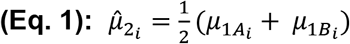

where 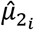 represents the predicted mean deflection, or safety margin, around the obstacle on 2-target trials for each participant *i*, and 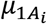 and 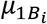 represent each participant’s observed mean deflection, or safety margin, on obstacle-obstructed and unobstructed 1-target trials, respectively. In Expt 2a, where movements on obstacle-obstructed left 1-target trials were deflected rightward, MA consequently predicts a rightward movement deflection on obstacle-present 2-target trials. In Expt 2b, where movements on obstacle-obstructed left 1-target trials were deflected leftward, MA consequently predicts a leftward movement deflection on obstacle-present 2-target trials (see Fig 3d).

Importantly, the obstacles in Expt 2a and 2b were designed not only to induce opposite deflections on obstacle-obstructed 1-target trials, but also to display identical protrusions in the direction of the intermediate movements that occur during uncertainty. Thus a motor plan optimizing task performance for intermediate movements during uncertainty would not be differentially affected by these obstacles. Specifically, a model for PO would have two objectives on obstacle-present 2-target trials: (1) to reach the final target within the required timing criteria and (2), to avoid obstacle collision, as these two objectives form the basis of task success (see methods). We therefore modeled the PO predictions as an average of the movement deflections that would arise if PO were to independently prioritize each objective, expressing the PO prediction as an equal balance of the two motor costs associated with the determinants of task performance. Prioritization of movement timing, or rapid target acquisition, would predict a movement direction midway between the two potential targets (i.e., a net 0° deflection), as this would maximize the probability of successful target acquisition during uncertainty^34^. However, prioritization of obstacle avoidance would predict deflections that incorporate a safety margin around the obstacle that is proportional to an internal estimate of variability^42^. To determine the expected safety margin for obstacle-present 2-target trials, we therefore scaled the magnitude of the safety margin observed on obstacle-obstructed 1-target trials by the ratio of their observed variabilities,

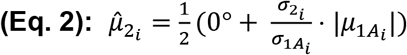

where 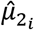 again represents the predicted mean deflection, or safety margin, on obstacle-present 2-target trials for each participant *i*, 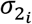 and 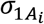 represent each participant’s observed variability on obstacle-present 2-target trials and obstacle-obstructed 1-target trials respectively, and 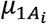 again represents each participant’s observed mean deflection, or safety margin, on obstacle-obstructed 1-target trials. The inclusion of this safety margin in motor planning for PO predicts deflections that skew *away* from the obstacle in both Expts 2a and 2b. Thus, the MA and PO hypotheses both predict rightward deflections that skew away from the obstacle on 2-target trials in Expt 2a, but in Expt 2b, MA predicts leftward deflections that skew towards the obstacle, whereas PO continues to predict rightward deflections that skew away from the obstacle.

### Performance-optimization predicts motor planning for obstacle avoidance

Like in Expt 1, we focused our analysis on the initial portion of motor output, measured as the initial movement direction (IMD), as this reflects feedforward motor planning. We calculated the IMD as the direction of the hand at the midpoint of the movement relative to the direction of the hand at movement onset (but note that we obtained qualitatively similar results for all analysis when the IMD was calculated 4cm into the movement). Each 2-target trial IMD was subsequently assigned a positive polarity if a deflection away from the obstacle occurred, and a negative polarity if a deflection towards the obstacle occurred. To facilitate a fair comparison between obstacle-free and obstacle-present 2-target trial IMDs (Fig 4b), the polarity of obstacle-free 2-target trials was arbitrarily assigned from one trial to the next. Correspondingly, on obstacle-present 2-target trials, we found that participants displayed IMDs that were consistently biased *away* from the obstacle compared to baseline obstacle-free IMDs in both Expt 2a (6.3 ± 4.3° vs −2.1 ± 2.4° [mean ± 95% CI], *p* = 7.90 × 10^−3^, t(7) = −3.16), and Expt 2b (3.6 ± 0.77° vs −0.2 ± 0.6°, *p* = 2.42 × 10^−4^, t(25) = −4.01; *paired t-test*). Example trajectory data is shown in Fig 4a, population-averaged IMD data in Fig 4b, and all individual participant IMD data in Fig 5b,e. The bias observed is consistent with the predictions of PO, but at odds with the predictions of MA. Moreover, we found that the observed IMDs were closer to the PO prediction than the MA prediction by a large margin in Expt 2b (see Fig 4b and Fig 5a,d), where gross differences in the PO and MA predictions were expected (root-mean-squared-errors (RMSEs) of 12.3 ± 1.5° and 2.4 ± 0.7° for MA and PO respectively; *p* = 1.22 × 10^−14^, t(25) = 15.9; two-sided *paired t-test*). Somewhat surprisingly, we also found that IMDs were closer to the PO rather than the MA prediction, albeit by a smaller margin in Expt 2a, where gross differences in the MA and PO predictions were not expected (RMSEs of 7.8 ± 1.9° and 2.78 ± 0.9° for MA and PO respectively; *p* = 7.10 × 10^−3^, t(7) = 3.75; *paired t-test*). Collectively, these findings suggest that during uncertainty, participants form a single action plan that optimizes task performance instead of automatically averaging potential motor plans.

### Refinement of the motor averaging and performance-optimization models

To allow the MA and PO models to make predictions without free parameters, we assumed equal weightings for both potential targets in the MA model (in line with previous work^1,2,15,20^), and analogously, for both determinants of the PO model (obstacle avoidance and rapid target acquisition). While this is reasonable, we assessed whether including differential weighting for the obstacle-obstructed and unobstructed targets might provide an explanation for why the original MA model could not accurately predict the experimental data. Analogously, we assessed whether including differential weighting for obstacle avoidance and rapid target acquisition might improve the already-accurate predictions from the PO model. Therefore, we evaluated refined versions of the original MA and PO models that each incorporated a single free weighting parameter (see methods) by using the Expt 2a data to fit that parameter, and the Expt 2b data to test the subsequent prediction for each model.

We found that the weighting parameter for the refined MA model that best fit the Expt 2a data produced weighting coefficients of 0.66 and 0.34 for the obstacle-obstructed and unobstructed targets, respectively. Correspondingly, we found that the best fit weighting parameter for the refined PO model produced weights of 0.29 and 0.71 for movement timing and obstacle avoidance, respectively. For the Expt 2a data, on which the weighting parameters were fit, the refined MA and PO models displayed similar RMSEs (3.07 ± 1.29° and 2.62 ± 0.97° for refined MA and refined PO RMSEs respectively, *p* = 0.46, t(7) = 0.78; *paired t-test*; see Fig 5a). However, for the Expt 2b data, the refined MA model resulted in predictions that were even worse than the original MA model (23.4 ± 1.8° vs 12.3 ± 1.5°). In contrast, the refined PO model resulted in predictions that were similar to the original PO model (3.8 ± 1.0° vs 2.4 ± 0.7°), consequently leading to an enhanced discrepancy between the refined MA and PO model predictions based on RMSEs (*p* = 1.01 × 10^−22^, t(25) = 34.8; *paired t-test*; see Fig 5d). The worsened prediction for the MA model produced by this weighting refinement indicates that the assumption for equal weighting present in the original model was not the reason that it failed to explain the experimental data.

### Performance-optimization predicts individual differences in obstacle avoidance

Panels 5b and 5e illustrate the relationship between the 2-target trial IMD, 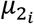, and the obstacle-obstructed 1-target trial IMD, 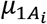, the latter of which was determined as the IMD relative to the angular protrusion of the obstacle into the direction of the observed deflection (i.e., the safety margin). We specifically plotted these variables because the MA and PO models both predict relationships between them (see Eq. 1–2), however, in both cases, the relationship between the predicted 2-target trial IMD, 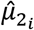, and 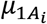 is not a singular one. In the MA model, 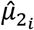 also depends on 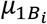, and in the PO model, 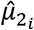 also depends on 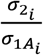. Correspondingly, these panels show the raw data of 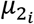 (black circles) as well as versions of 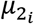 that correct for the effects of 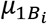 (purple circles) to evaluate the MA model predictions, and for the effects of 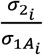 (green circles) to evaluate the PO model predictions. We found that 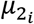, when corrected for 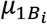, did not display a consistent positive correlation with 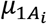, as the correlation was positive for the data from Expt 2a (*r* = +0.91, *p* = 1.51 × 10^−3^, t(6) = 5.49; 1-sample *t-test*) but not for the data from Expt 2b (*r* = −0.22, *p* = 0.28, t(24) = −1.11; 1-sample *t-test*). In contrast, we found that 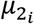, when corrected for 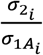, consistently displayed a positive correlation with 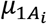 (Expt 2a: *r* = +0.88, *p* = 3.70 × 10^−3^, t(6) = 4.60; Expt 2b: *r* = +0.71, *p* = 4.89 × 10^−9^, t(24) = 4.94; 1-sample *t-test*). This suggests that performance-optimization can explain not only mean behavior during obstacle avoidance, but also differences between one individual and another.

To better understand the ability of the PO model to predict the individual differences present in 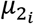, we next sought to rigorously examine the importance of individuating each predictor, 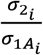 and 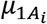. To accomplish this, we devised an extension of the PO model that attempted to explain 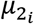 using a combination of individuated and non-individuated versions of each predictor,

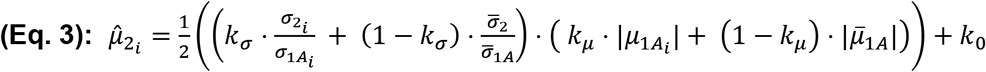

which includes three free parameters in *k*_*σ*_, *k*_*μ*_, and *k*_0_ (an offset term). Here a value of 1 for *k*_*σ*_ would indicate that individuating the variability ratio 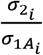 is far superior to using the population-averaged ratio 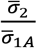 in explaining 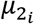, whereas a value of 0 would indicate the reverse. Analogously, a value of 1 for *k*_*μ*_ would indicate that individuating the obstacle-obstructed 1-target trial IMD 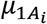 is far superior to using the population-averaged IMD 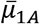 in explaining 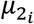, whereas a value of 0 would indicate the reverse. We fit this model onto the pooled aggregate of datasets from Expt 2a and 2b, and found that it was able to explain 79.5% of the variance for individual differences in 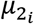, with values of 0.71 and 0.63 for *k*_*σ*_ and *k*_*μ*_, respectively, indicating that 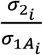 and 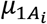 both contribute substantially to the ability to predict individual differences in the 2-target trial IMD 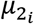. Correspondingly, we found similar contributions to the explained variance from 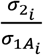 and 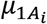 (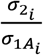: partial *R*^2^ = 0.63, *p* = 5.52 × 10^−8^, *F*(1,31) = 50.5; 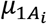: partial *R*^2^ = 0.46, *p* = 1.15 × 10^−5^, *F*(1,31) = 27.1; *F-test*).

Further dissection of the PO model, in which the obstacle-obstructed 1-target trial IMD 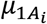 is expressed as the product of the obstacle-obstructed 1-target trial variability 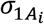 and a corresponding variability responsiveness factor 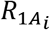, yields a model where 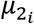 is predicted from 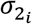 and 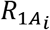,

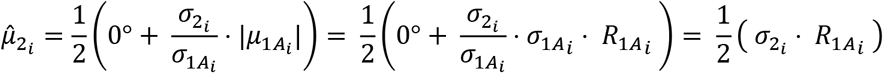

and parametrically individuating this equation yields,

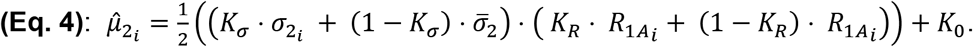

Here the parameters *K*_*σ*_ and *K*_*R*_ are analogous to *k*_*σ*_ and *k*_*μ*_ from Eq. 3. The variability responsiveness factor, 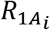, is a value that scales the safety margin, and thus corresponds to individual differences in confidence against obstacle collision. This version of the PO model explained 91% of the variance for individual differences in 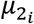, with values of 0.36 and 0.10 for *K*_*σ*_ and *K*_*R*_, respectively, indicating that 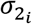 and 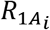 both contribute to the ability to predict individual differences in the 2-target trial IMD 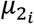. However, whereas 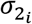 explained a large fraction of variance in 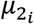, 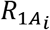 explained much less (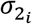: partial *R*^2^ = 0.90, *p* = 1.01 × 10^−26^, *F*(1,31) = 314; 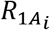: partial *R*^2^ = 0.19, *p* = 0.012, *F*(1,31) = 7.2; *F-test*). These results suggest that motor variability drives the PO model’s ability to predict the inter-individual differences observed in Expt 2.

Figs 5c and 5f show the relationship between inter-individual differences in 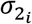 and inter-individual differences in 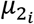 in Expt 2a and 2b respectively. We exploited this relationship to assess additional predictions for MA and PO. A safety margin dependent PO hypothesis would predict a significantly positive slope between these variables, but because MA determines IMDs during uncertainty based entirely on the mean of obstacle-obstructed 1-target trial IMDs 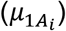, it predicts a null, or zero-slope relationship. In line with the modeling results presented above, both Expts 2a and 2b exhibited strong positive correlations between *σ*_2_ and *μ*_2_ (Expt 2a: *r* = +0.96, *p* = 1.51 × 10^−4^, t(6) = 8.49, Fig 5c; Expt 2b: *r* = +0.89, *p* = 2.60 × 10^−9^, t(24) = 9.16; 1-sample *t-test*; Fig 5f) and indicate that *μ*_2_ is approximately linearly sensitive to *σ*_2_ (Expt 2a: slope = 0.81, *R*^*2*^ = 0.78; Expt 2b: slope = 0.94, *R*^*2*^ = 0.93), in agreement with previous work investigating the relationship between the safety margin for actions and variability^42^. The participant with the highest measurement for 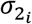 in Expt 2a, and the two participants with the highest measurements for 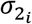 in Expt 2b, epitomized this relationship by correspondingly demonstrating the highest levels of 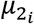. However, even when we excluded these participants, positive correlations remained (Expt 2a: *r* = +0.80, *p* = 0.032, t(5) = 2.95; Expt 2b: *r* = +0.73, *p* = 1.02 × 10^−4^, t(22) = 4.87; 1-sample *t-test*). Collectively, these results indicate that safety margins during uncertainty are highly sensitive to directional variability, in line with a PO hypothesis, echoing both the population-averaged and individual-participant based modeling predictions, as well as the findings from Expt 1.

## DISCUSSION

We designed two novel experimental paradigms that powerfully dissociated between the hypotheses proposed to underly motor planning under uncertainty: motor averaging (MA) and performance-optimization (PO). In Expt 1, we designed an environment that physically perturbed motion in the direction of potential target locations off-course, and motion in the direction of intermediate movements oppositely off-course. Participants readily adapted to this composite environment on 1-target trials. Critically, on trials with two potential targets, participants displayed motor output strongly aligned with adaptive responses for intermediate movements, which was consistent with the PO prediction, but grossly opposite the MA prediction for motor planning under uncertainty (see Fig 2). An effort to refine the model predictions by taking the observed spread of movement directions into account, resulted in even greater alignment between the observed motor output and the PO prediction.

In Expts 2a-b, we replicated (Exp 2a) the paradigm from a well-known study that provided support for MA^8^ but did not qualitatively dissociate the predictions for MA and PO, and then made a small modification (Exp 2b) that allowed for MA and PO to be powerfully dissociated. Specifically, in Expt 2a, we positioned an obstacle so that movements to one of the potential targets would be skewed in a direction consistent with increasing the safety margin for intermediate movements during uncertainty. Thus, qualitatively, the PO prediction based on creating an appropriate safety margin for intermediate movements during uncertainty, and the MA prediction based on averaging the motor plans for potential targets, were in the same direction. But quantitatively, we found the experimental results to be significantly closer to the PO predictions than the MA predictions. In Expt 2b, we altered the obstacle from Expt 2a so that movements to one of the potential targets would be skewed in a direction opposite of that needed for increasing the safety margin for intermediate movements during uncertainty, resulting in both qualitative and quantitative differences in the predictions for MA and PO. Experimentally, we found that motor output during trials with two potential targets consistently increased the safety margin for intermediate movements in accordance with the PO prediction, but was grossly opposite in direction to obstacle-induced changes in the MA prediction. Subsequent refinement of the MA model to allow for different weightings of the motor plans associated with the obstacle-obstructed and unobstructed targets did not improve its prediction. On the other hand, the PO model accurately predicted the population averaged changes in motor output for both experiments as well as a remarkable ~80% of the variance for individual differences between participants, suggesting that internal estimates of variability and uncertainty can be critical for motor planning. Collectively, our results provide clear evidence that humans generate a single motor plan that reflects optimization of task performance, rather than averaging over multiple potential plans.

### Previous work suggesting motor averaging for motor planning under uncertainty

Our finding that performance-optimization underlies movements executed during uncertainty challenges the long-standing idea that the intermediate movements executed during uncertain conditions reflect motor averaging. However, many studies suggesting that either sensory or motor representations of movement plans are averaged did not dissociate these possibilities from selecting a single plan that optimizes motor performance. For example, the Stewart et al. 2014 study^8^ that we replicate in Expt 2a, attempted to dissociate whether motor output might reflect an average of sensory or motor representations of movement plans (i.e., sensory averaging vs motor averaging) by using obstacles to perturb these motor plans, but did not dissociate either sensory or motor averaging from performance-optimization.

Another study, by Gallivan et al. (2017)^16^, applied a visuomotor rotation (VMR) to movements toward one potential target to perturb motor planning without affecting the sensory representation of the target. This resulted in initial movement directions (IMDs) that were altered during 2-target trials in accordance with the perturbed motor plans, and thus in line with the prediction for motor, rather than sensory, averaging. However, PO predicts an IMD identical to that predicted by MA: Since the imposed VMR shifts the final hand position associated with acquisition of the potential target to which it was applied, a corresponding shift in the IMD that accounts for this VMR perturbation would optimize performance by minimizing the cost of corrective movements following disclosure of the final target location. Therefore, this experimental manipulation, like that in Stewart et al. (2014) and a number of other studies^1,8,12,15,17–20^, fails to dissociate MA from PO.

### Previous work on performance-optimization for motor planning under uncertainty

The challenge of examining motor planning under uncertainty has persisted even for studies that made deliberate attempts at dissociating MA from PO. For example, in one study performed by Wong and Haith in 2017^9^, motor planning under uncertainty was examined in conditions where participants were required to reach towards one of two potential targets with a movement completion time that was small enough to preclude movements with corrective adjustments from being successful. The results showed that under these conditions, participants frequently abandoned intermediate movements that subsequently require corrective adjustments on 2-target trials in favor of direct movements towards one of the potential targets. This finding was interpreted to indicate that PO, rather than MA, underlies motor planning under uncertainty. However, it is likely that participants simply made a strategic decision to guess at the final target location and aim at that guess. While this would indeed improve performance and could therefore be considered a type of performance-optimization, such strategic decision making does not provide information about the implicit neural processing involved in programming the motor output for the intermediate movements that are normally planned under uncertain conditions.

In another study, performed by Nashed and colleagues (2017)^35^, motor planning under uncertainty was studied using a task that was based on grip force (GF) control. Specifically, participants grasped an object, capable of measuring GF, and made reaching movements while environmental dynamics were experimentally manipulated with robotically applied load forces that affected the required GF. This task design was used to attempt for a dissociation of MA from PO, however, although GF is known to be substantially more sensitive to the variability in environmental dynamics than to the mean dynamics^42^, this study examined the MA and PO hypotheses for motor planning under uncertainty using predictions that were based entirely on the mean environmental dynamics. Taken together with the lack of information provided about environmental variability and its effect on required GFs, it is unlikely that this study can shed light on motor planning under uncertainty.

### Neural representations of motor planning under uncertainty

Studies suggesting motor or sensory averaging have been motivated by reports of simultaneous deliberation of competing potential goals in sensorimotor brain areas (i.e., parallel planning)^1,8,12,18,33^. These studies argue that the motor plans prepared in parallel are averaged, resulting in the intermediate movements observed when uncertainty is present during motor planning. The neural evidence for parallel planning is based on studies of delay period activity associated with motor planning when multiple potential targets were present^28,29,31,32^. However, because this activity was measured using single-electrode recordings, only a small number of cells could be recorded simultaneously, and thus data from same-type trials were aggregated to make population-based estimates of the planned motion. The results suggested that this aggregate delay period activity was tuned to both potential targets in sensorimotor areas, in particular dorsal premotor cortex (PMd) and parietal reach region (PRR), in monkeys. However, the observed tuning could arise from simultaneous parallel planning for both potential targets, as the authors argued, or from planning-related activity associated with one target on some trials and with the alternate target on other trials.

Recently, Dekleva and colleagues (2018)^43^ tested for parallel planning using an electrode array to record simultaneously from 100-160 neurons in PMd so that planning-related neural activity could be examined for individual movements when two potential targets were present. Critically, they found that neural activity during the delay period was consistent with motor planning directed for either one target or the other on individual trials, rather than with parallel planning for both potential targets. While these results cannot rule out parallel planning in other brain areas, they call into question the evidence for parallel planning from previous studies that relied on trial-aggregated data.

In summary, the current findings indicate that motor planning during uncertain conditions does not proceed from averaging parallel motor plans, but instead, incurs the creation of a single motor plan that optimizes task performance given knowledge of the current environment. These findings are compatible with the current neurophysiological data and offer a mechanistic framework for understanding motor planning in the nervous system.

## MATERIALS AND METHODS

### Participants

Twenty-six participants (twenty-five right-handed; 15 female; age range 18-42) performed the multi-force-field (multi-FF) adaptation experiments, with sixteen participants in Expt 1 and ten participants in Expt 1-GEN. Thirty-four participants (thirty-two right-handed; 18 female; age range 18-33) performed the obstacle avoidance experiments, with eight participants in Expt 2a and twenty-six participants in Expt 2b. Participants were assigned to experiments based when they responded to advertisements their availability for scheduling. When different experiments were running concurrently, participants were randomly assigned for participation. The sample sizes used for each experiment were determined based on pilot data and existing literature^8,38,39^. All participants used their right hands to perform the experiments, were naïve to the purpose of the experiments, and were without known neurological impairment. The study protocol was approved by the Harvard University Institutional Review Board, and all participants provided written informed consent.

### Experiment Protocols

#### Apparatus for multi-force-field adaptation experiments (Expt 1 and Expt 1-GEN)

Participants were instructed to grasp the handle of a two-joint robotic manipulandum with their right hands and make rapid 20cm point-to-point reaching arm movements in the horizontal plane to either a single pre-specified target (1-target trials) or two potential targets (2-target trials). All visual information, including veridical feedback of hand motion provided in the form of a white 3mm-diameter cursor, was displayed on a vertically oriented LCD computer monitor (refresh rate of 75 Hz). Participants were positioned such that their midlines were aligned with the middle of the monitor, and their right arms were always supported with a ceiling-mounted sling. The manipulandum measured hand position, velocity, and force, and its motors were used to dynamically apply prescribed force patterns to the hand, all of which were updated at a sampling rate of 200Hz.

#### Targets and feedback (Expt 1 and Expt 1-GEN)

On 1-target trials, the target was located at a left (+30°), right (−30°), or center (0°) eccentricity from the midline, and on 2-target trials, a pair of potential targets always appeared at the left and right target locations. Participants were instructed to initiate a trial by moving the cursor to a start position (green filled-in circle, 10mm-diameter), after which the target or targets (yellow hollow circles, 10mm-diameter) were presented. 1000ms after target presentation, an auditory go cue signaled participants to initiate a movement. Movement onset was subsequently determined online as the time when the hand velocity exceeded 5cm/s or the time when the hand traveled 3cm from the start position, whichever occurred first. Participants were required to initiate movements after the onset of the go cue, but no later than 425ms after the go cue finished playing. If movement onset was detected outside these bounds, a message that either read ‘Too Soon!’ or ‘Too Late!’ was displayed above the start position, and was accompanied with a unique tone. Furthermore, because pilot data showed that participants may sporadically stop and initiate a discrete reach to the final target immediately following its disclosure on 2-target trials, we also required participants to maintain their velocity throughout the first half of every movement (i.e., while the displacement was less than 10cm). Specifically, during each trial, if the instantaneous maximum velocity declined more than 33% during the first half of the movement, we played a unique buzzer tone. If any of these requirements were not fulfilled, the trial was discarded.

For the 2-target go-before-you-know trials, the final target (randomly selected on each trial) was filled in with yellow, and the distractor target was simultaneously extinguished once a 3cm displacement between the hand and start position was achieved. For consistency, on 1-target trials, the target also filled in with yellow at the same point in the movement. After the hand reached the final target, we provided performance feedback based on the movement time, determined as the time interval between movement onset, as defined above, and movement offset, defined as the first timepoint when the hand was both within 6mm of the final target, and the hand speed in the subsequent 300ms period was below a threshold of 6.35cm/s. Following movement offset, visual and auditory feedback were presented by changing the fill color of the target and playing a sound, depending on whether the movement time was faster than (red fill-in color, buzzer tone), within (green fill-in color, chirp tone), or slower than (blue fill-in color, buzzer tone) the required interval. This movement time interval was based on thresholds that were adjusted online per participant as described below. After feedback was delivered, the robotic manipulandum guided participants’ hands back to the start position. Participants completed blocks of trials throughout the experiments but were allotted 1min rest breaks in-between each block (see training schedule details below).

#### Movement time thresholds

We used data-driven updating for specifying the movement time thresholds. The “too-slow” threshold was set at the 70^th^ percentile of the of the movement times for last 18 trials of the same type (1-target or 2-target). Thus, separate thresholds were maintained for 1-target vs 2-target trials. The 1-target trial movement time threshold ranged from 225 to 585ms in Expt 1 and 225 to 608ms in Expt 1-GEN (on average across participants). In Expt 1, the 2-target trial movement time thresholds ranged from 225 to 655ms. We used this 70^th^ percentile updating to individualize the movement time thresholds for different participants because we found that, in pilot data, individuals with a large fraction of “too-slow” feedback sometimes exhibited erratic, seemingly exploratory behavior on 2-target trials, and abandoned intermediate movements. Individuals with a smaller fraction of “too-slow” feedback, however, did not. Intermediate movements are of fundamental interest for the examination of motor planning during uncertainty as these movements reflect low-level implicit motor planning^16^, but unfortunately, abandonment of intermediate movements has remained an issue in studies with standard implementations of go-before-you-know tasks, consequently leading to striking data exclusion criteria (e.g., removal of 7%-50% of participants in previous work^5, 12–14, 25–26^). Individualized movement time thresholding allowed us to evoke intermediate movement behavior consistently within and across all participants.

#### Multi-force-field environment

To dissociate between performance-optimization (PO) of intermediate movements during goal uncertainty and motor averaging (MA) of actions associated with each potential goal presented during uncertainty, we differentially perturbed the 1-target motor plans in a manner that resulted in distinct predictions for each hypothesis in Expt 1. To create such a perturbation, we used the robot motors to produce a dynamic environment comprised of multiple curl force-fields (FFs) that acted on the manipulandum to perturb hand motion off-course in opposite directions depending on the cued target direction during 1-target trials in Expt 1 and Expt 1-GEN. This multi-FF environment levied velocity-dependent FFs that were proportional in magnitude and directionally orthogonal to the velocity of hand motion,

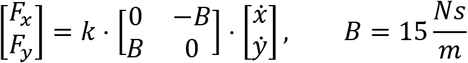

where 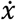 and 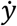 denote the hand velocities, 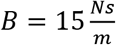 denotes the velocity-dependent gain, and *k* = ±1 denotes a binary switch variable that was set to opposite values for center target trials versus left target or right target trials for each participant. Setting *k* = +1 results in clockwise (CW) FFs whereas *k* = −1 results in counterclockwise (CCW) FFs. We balanced the directions of the applied FFs across participants so that half experienced the multi-FF environment with *k* = +1 for left target or right target trials (e.g., see FF_A_ in Fig 1c) and *k* = −1 for center target trials (e.g., see FF_B_ in Fig 1c), and the other half experienced the multi-FF environment with *k* = −1 for left target or right target trials and *k* = +1 center target trials. Data associated with each target were then combined from each subgroup.

#### Error clamp and partial error clamp trials

Because actions made during reaching movements may result from both feedforward motor planning or online feedback corrections to movement errors, we used error clamp (EC) trials to restrict deviations from the straight-line path towards the target during 1-target trials. We implemented these EC trials as a highly stiff (6000 N/m), viscous (250 Ns/m) one-dimensional spring and damper system in the direction orthogonal to the straight-line path between the initial hand position and the cued target. In line with previous work, these EC trials effectively eliminated movement errors (average maximum absolute deviation, <1.9mm), and allowed for a high-accuracy measurement of feedforward participant-produced forces patterns^37,39^.

Unlike 1-target trials, feedback corrections are to be expected during 2-target trials if task success is to be achieved, because participants must reflexively correct their movements following divulgence of the ultimate target. We thus devised a variant of the EC, which we term the partial error clamp (pEC), to measure the initial segment of force output during 2-target trials that reflects feedforward motor planning *before* these feedback corrections occur. As the initial motion on these 2-target trials was directed towards the center target, we aligned the pECs to the straight-line path to the center target, but we smoothly transitioned the movement from the highly stiff and viscous environment into a null environment (i.e., the robot motors were disabled) after 11cm. Note that the 11cm point was selected based on an analysis of pilot data to determine the onset of feedback corrections to the final target. These pEC trials allowed us to measure of feedforward force patterns early in the movement, while motor errors were minimized (average maximum absolute deviation, <2.3mm), but still permitted participants to carry out feedback corrections later in the movement for final target acquisition. Post-hoc surveys indicated that 5/16 participants noticed pECs, whereas 6/16 participants noticed the ECs.

#### Training schedules (Expt 1 and Expt 1-GEN)

We divided the experiment into baseline, training, and test epochs with a total of 1305 outward reaching movements. The experiment began with the baseline epoch, which consisted of nine blocks. The first three blocks were comprised of 120 null 1-target trials that familiarized participants with the basic experimental setup and feedback structure described above. The next four blocks were comprised of 200 2-target trials, the first 50 of which were null trials, and the remaining 150 were 80% null trials and 20% pEC trials. The last three blocks of the baseline period reacquainted participants with 1-target trials before the training period started, and were comprised of 85 1-target trials, in which 80% were null trials and 20% were EC trials. The force patterns measured on EC and pEC trials throughout this epoch were used as a baseline for estimating learning-related changes in force patterns during subsequent blocks. Note that in this epoch, all blocks comprised of 1-target trials probed each target direction in equal amounts.

The baseline epoch was followed by the training epoch, which was separated into seven blocks and was comprised exclusively of 1-target trials. The purpose of this epoch was to elicit robust adaptation to the multi-FF environment we designed. However, because this environment differentially perturbs movements to the center target compared to the lateral (left/right) targets (see *Multi-force-field environment*), interference effects for adaptation to the FFs associated with the lateral targets would arise from center target training. Importantly, whereas the source of interference for left/right target training is singular, center target training would suffer interference effects from the FFs associated with both lateral targets. Interference effects would limit the overall level of adaptation that can be achieved, and because these effects are non-uniformly distributed across the trained target directions, straightforward application of the multi-FF environment may lead to dissimilar levels of adaptation across target directions. These issues would consequently limit the ability to dissociate MA from PO and would bias their predictions. We therefore sought to create a training schedule that would avoid this bias and elicit similar levels of adaptation to the FFs associated with each target. To do so, we tested various training schedules in a pilot study that was guided by simulations from a linear state-space model with local motor primitives for predicting the effects of generalization in our composite environment. We correspondingly employed the FF training schedule we found successful, which included 500 training trials (with 80% FF trials and 20% EC trials), and twice the number of center target trials compared to the number of left or right target trials. Note that this distribution of target directions (1:2:1 for left, center, and right target directions, respectively) carried into the next epoch.

After the training epoch, participants completed the test epoch, which was separated into five blocks. The purpose of this epoch was to probe participants’ initial force patterns during 2-target trials while approximately maintaining the level of adaptation to the multi-FF environment achieved for 1-target trials during the training epoch. We thus included 400 trials in this epoch, of which 300 were 1-target FF trials, 50 were 1-target EC trials, and 50 were 2-target pEC trials. Note that during all epochs, ECs/pECs were interspersed in a pattern that was random (frequency of 1 in 5 during baseline and training and 1 in 4 during test) but which avoided consecutive EC/pEC trials to prevent decay.

The training schedule of Expt 1-GEN was analogous to that of Expt 1, but 2-target trials were replaced with 1-target trials that were positioned at 1 of 9 different directions (from −30° to 30° every 7.5°). Thus the training epoch of Expt 1-GEN was identical to that of Expt 1, but during the baseline epoch, participants reached towards each of the nine targets, presented in random order, for an equal number of trials. Baseline force patterns associated with each target were probed with ECs on 20% of trials after the third baseline block as in Expt 1. During the test epoch, the 2-target pEC trials from Expt 1 were replaced with 1-target EC trials that probed generalization to the targets that were positioned in-between the trained target directions (i.e., these trials probed the targets located at ±22.5°, ±15°, and ±7.5°).

#### Apparatus for obstacle avoidance experiments (Expt 2a and Expt 2b)

Participants were instructed to grasp a lightweight plastic handle that sheathed a digital stylus and make reaching movements with their right hands in the horizontal plane. We instructed participants to slide the handle across the surface of a tablet capable of recording hand position at 200Hz with a resolution of 0.01mm. All visual stimuli, including targets, obstacles, and a real-time cursor showing hand position, were displayed on a horizontally oriented LCD computer monitor (with a screen refresh rate of 120Hz and a motion display latency of ~25ms) that was mounted above the tablet at the shoulder level and therefore obstructed view of the hand. Participants were positioned such that their midlines were aligned with the middle of the monitor and tablet.

#### Design of obstacles (Expt 2a and Expt 2b)

In Expts 2a and 2b, participants made reaching movements using the 1-target and 2-target trial configurations from Expt 1, but on some trials, we presented visual obstacles that we instructed participants to avoid. As illustrated in Fig 3c, the obstacle in Expt 2a was rectangular in shape (width 1cm and length 12cm) and was oriented so that its long axis was perpendicular to the vector between the start position and the target. Moreover, it was positioned midway between the start location and the target, and from this location, protruded 2cm towards the midline and 10cm away from it so that movements around the obstacle would be consistently deflected towards the midline. In Expt 2b, we modified the obstacle from Expt 2a by clipping off the 10cm away-from-midline protrusion, so that it still protruded 2cm towards the midline, but now 0cm away from it (see Fig. 3c, bottom panel). This modification promoted deflections that were away from the midline, opposite in direction to those in Expt 2a, and allowed us to dissociate the MA and PO hypotheses for motor planning under uncertainty.

#### Trial types and feedback (Expt 2a and Expt 2b)

Expts 2a-b included obstacle-free and obstacle-present 1- and 2-target trials. Visual stimuli and feedback were identical to those of Expt 1. Participants were instructed to rapidly reach the final target but while avoiding a virtual obstacle, if present. If a collision with the obstacle was detected, a message that read ‘You hit the obstacle!’, was displayed, a buzzer tone was played, and the trial was disqualified from movement time reward feedback. Otherwise, the movement time feedback and the individualized thresholding procedure was identical to Expt 1, but with separate thresholds maintained for 2-target trials, obstacle-obstructed 1-target trials, and all remaining 1-target trials (i.e., all obstacle-free 1-target trials and all obstacle-present 1-target trials in which the obstacle did not directly block the target). We did not maintain separate thresholds for obstacle-free vs obstacle-present 2-target trials because pilot studies indicated that the differences in movement completion times were small. The 2-target trial movement time thresholds ranged from 225 to 645ms in Expt 2a and 225 to 598ms in Expt 2b (on average across participants). The obstacle-obstructed 1-target trial movement time threshold ranged from 225 to 450ms in Expt 2a and 225 to 360ms in Expt 2b. The movement time threshold for the remaining 1-target trials ranged from 225 to 380ms in Expt 2a and 225 to 382ms in Expt 2b. After participants reached the final target, they were instructed to move the handle back to the starting position to begin the next trial.

Note that all participants in both experiments completed an equal number of trials in which the obstacle was positioned between the start target and the left target (left-side obstacle condition) and between the start target and right target (right-side obstacle condition). Experiments were, therefore, balanced within participants to cancel out any target-specific effects that might lead to biases in movement direction. Fig 3 displays the MA and PO predictions based on a left-side obstacle condition, but for a right-side obstacle condition, the geometry of the predictions would simply be left/right mirror-reversed. Data shown in Fig 4b and Fig 5 correspondingly combine the left-side and right-side obstacle conditions for each participant.

#### Training schedules (Expt 2a and Expt 2b)

Expt 2a and 2b had identical training schedules and thus the experiments only differed in the obstacle geometry, as described above and shown in Fig 3. Both experiments were divided into a baseline and test epoch with a total of 760 outward reaching movements. The baseline epoch included nine blocks (460 trials total), and was designed to familiarize participants with the basic task and feedback structure, before obstacle-present 2-target trials were presented in the test epoch. The first two blocks were comprised of 120 obstacle-free 1-target trials, and the next two blocks were comprised of 100 obstacle-present 1-target trials. Of the five blocks that followed, which included a total of 240 trials, three were comprised of obstacle-free 1- and 2-target trials (with 40% 1-target trials and 60% 2-target trials, 180 trials total) and the remaining two blocks were comprised solely of obstacle-present 1-target trials (60 trials total). Note that across all baseline epoch trials, 1-target trials probed each target direction in equal amounts.

The test epoch included six blocks (300 trials total), and was designed to probe motor planning for obstacle-present 2-target trials so that predictions for MA and PO could be compared. All blocks in this epoch were comprised solely of obstacle-present trials, with 60% obstacle-present 1-target trials and 40% obstacle-present 2-target trials. Of the obstacle-present 1-target trials, we probed movements towards the obstacle-obstructed target more often (50% of trials) because our analyses were more sensitive to movements towards this target compared to movements towards the center or unobstructed targets. The remaining 50% of obstacle-present 1-target trials probed the center and unobstructed targets in equal amounts. Note that, in both the baseline and test epochs, left-side and right-side obstacle conditions were separated into different blocks, and the ordering of these blocks was balanced across participants.

### Analysis

#### Outlier Analysis

No participants were excluded from any dataset. Individual movements that did not comply with the task requirements, outlined in *Targets and feedback (Expt 1 and Expt 1-GEN)*, were not eligible for analysis (<3% of trials in Expt 1, <2% of trials in Expt 1-GEN, <1% of trials in Expt 2a, and <2% of trials in Expt 2b). In addition, we discarded a small fraction of highly atypical movements based on two key features. For all experiments, we required that the movement time was between 225ms and 2000ms (<2% of trials in Expt 1, <1% of trials in Expt 1-GEN, <1% of trials in Expt 2a, and <1% of trials in Expt 2b) and for Expt 1 and Expt 1-GEN, we also required that peak velocity was between 0.2m/s and 1m/s (<1% of trials in Expt 1 and <1% of trials in Expt 1-GEN). Note that only movements performed after the familiarization blocks in each experiment were used for analysis, and for those movements, these criteria collectively resulted in the omission of <3% of trials in Expt 1, <2% of trials in Expt 1-GEN, <1% of trials in Expt 2a, and <2% of trials in Expt 2b.

#### Analysis of force patterns in Expt 1

We examined the lateral force profiles participants produced that were orthogonal to the cued target direction for 1-target EC trials and to the center target direction for 2-target pEC trials, corresponding to the axis of the imposed perturbations (see *Multi-force-field environment*). We aligned all force profiles to the onset of the target cue (T_ON_, 40-50ms after movement onset), and used the population-averaged force profiles measured during the test period, after participants became acclimated to the mutli-FF environment, to construct the MA and PO predictions. Specifically, we constructed the MA prediction by averaging the force profiles associated with the left and right targets, and we constructed the PO prediction by directly using the force profiles associated with the center target (Fig 2c). Since we sought to isolate the feedforward component of the data and predictions, before feedback responses to the target cue occurred, we analyzed all force profiles until the minimum time (across participants) that differences in force output on left and right cued 2-target pEC trials were significantly different from zero (T_RESP_, ~150ms after T_ON_). In addition, because the imposed FF environment was velocity-dependent, and adaptive responses to velocity-dependent dynamics are known to be scaled by movement velocity from one trial to the next^44^, we normalized each force profile by the velocity-dependent level of ideal compensation.

For a simple determination of how participants compensated for the multi-FF environment we imposed, we characterized the adaptive response on 1-target trials with an adaptation coefficient (AC), calculated as the slope from a linear regression of the baseline-subtracted force profiles participants made during EC trials onto the ideal compensatory force^38,39,42^. For trials that were associated with the FF perturbations imposed during movements towards the center target, we defined the AC so that full FF compensation would yield an AC of +1. For trials that were associated with the FF perturbations imposed during movements towards the left or right target, we defined the AC so that full FF compensation would yield an AC of −1.

To quantify the similarity between the 2-target trial force data and the predictions, as shown in Fig 2g, we devised a prediction index (PI) that results in a value of +1 if the 2-target trial data is perfectly similar to the PO prediction, −1 if it is perfectly similar to the MA prediction, and 0 would if the data were halfway between both predictions,

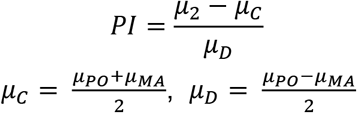

where *μ*_*C*_ and *μ*_*D*_ correspond to the common and differential modes, respectively, of predicted mean force levels based on the PO (*μ*_*PO*_) and MA (*μ*_*MA*_) predictions, and *μ*_2_ corresponds to the mean force level of the 2-target trial data. We calculated the PI over two intervals: one spanned movement onset until T_ON_, and the other spanned movement onset until T_RESP_.

#### Refinement of predictions based on generalization of adaptive responses (Expt 1 and Expt 1-GEN)

Due to non-trivial variability in motor output, participants occasionally deviated from the intended target direction on 1-target trials and from the center target direction on 2-target trials. Directional deviations consequently bias both the MA and PO predictions since adjacent targets were associated with different FFs in the composite environment we designed. To account for this variability-induced effect and refine our predictions, we first measured how the multi-FF environment generalizes to nine different movement directions in Expt 1-GEN (see *Training schedules (Expt 1 and Expt 1-GEN)*). We then estimated the generalization of adaptation throughout our composite environment by fitting the population-averaged ACs (from the test period) associated with every probed target direction onto a model that was based on the additive combination of Gaussians centered around the trained target directions (+30°/0°/−30°),

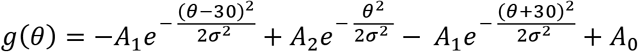

This equation describes the amount of generalization, *g*, as a function of the probed target direction, *θ*. There are four free parameters: *σ* is the width of each Gaussian, *A*_1_ and *A*_2_ are the heights of the Gaussians associated with left/right target training and center target training, respectively, and *A*_0_ is an offset. This model uses equal width Gaussians for the adaptation at each target, which is reasonable since FFs levied in each target direction were structurally identical. We allowed different heights to be associated with the left/right vs center target directions since movements towards the center target were perturbed twice as many times as movements towards the left or right targets (see *Training schedules (Expt 1 and Expt 1-GEN*)).

We used the model for *g*(*θ*) to determine how the force profiles associated with the MA and PO predictions from Expt 1 would be refined given each participant’s distribution of movement directions. We determined the movement direction as the direction of the hand when it was 10cm away from the start position relative to the direction of the hand at movement onset. Note that the distributions of movement directions shown in Fig 2e were based on random samples from the pooled aggregate of participant data. We used each participant’s distribution of movement directions on left and right 1-target FF trials (from the test epoch) to refine the MA predictions, and the movement directions on null 2-target trials to refine the PO predictions. We applied the model for *g*(*θ*) to predict the pattern of generalization up to ±42.5° away from the center target direction to obtain an estimate of generalization that encompassed the space of movement directions explored by participants. We then convolved the resulting model-estimated generalization function with each participant’s distribution of movement directions from Expt 1 to determine the corresponding expected value for the adaptative responses conditioned on the observed directional variability. Each participant’s expected level of adaptation was then used to scale the force profiles that previously formed what we refer to as the raw MA and PO predictions, and ultimately determined what we refer to as the refined predictions plotted in Fig 2f.

#### Motor averaging and performance-optimization models in Expt 2a and Expt 2b

As outlined in the results, the MA model posits that on 2-target trials, where uncertainty about the final goal location is present, individuals exhibit a motor plan that reflects an average of the motor plans associated with each potential goal,

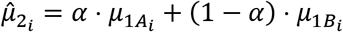

where 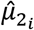 represents the predicted mean deflection, or safety margin, around the obstacle on 2-target trials for each participant *i*, and 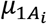 and 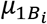 represent each participant’s observed mean IMD deflection on obstacle-obstructed and unobstructed 1-target trials, respectively. Here *α* is a weighting parameter that controls the influence of the motor plans associated with the obstacle-obstructed and unobstructed target. Thus *α* = 1 indicates that participants assigned all weight to the motor plan associated with the obstacle-obstructed target and *α* = 0 indicates that participants assigned all weight to the motor plan associated with the unobstructed target. In line with canonical MA theories^1,2,15,20^, we assumed 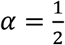 for the original MA model presented in Eq. 2 and for the corresponding predictions presented in Fig 5b,e.

The PO model posits that on 2-target trials, individuals exhibit a motor plan that attempts to achieve task success given knowledge of the environment. Thus, on obstacle-present 2-target trials, PO of intermediate movements would have two objectives: (1) to reach the final target within the required timing criteria and (2), to avoid obstacle collision, because as outlined in *Trial types and feedback (Expt 2a and Expt 2b)*, these two objectives determined reward on each obstacle-present trial. We modeled the PO prediction as a combination of the predicted IMDs that would arise if PO were to independently optimize each objective. Optimization of (1), movement timing, for task performance would lead to IMDs in the center target direction (0°), as this IMD maximizes the probability of successful target acquisition during uncertainty^5,34^. On the other hand, to determine the IMD predicted if individuals were to optimize (2), obstacle avoidance, we exploited the observation that the motor system linearly modulates the size of safety margins for actions based on internal estimates of motor variability^42^. Accordingly, we determined the expected safety margin around the obstacle during uncertainty by scaling the magnitude of safety margins observed during goal *certainty* by the increase in variability that is induced by goal *uncertainty*. Thus the PO model took the following form:

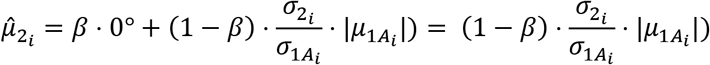

where 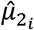 again represents the predicted mean deflection, or safety margin, on obstacle-present 2-target trials for each participant *i*, 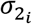 and 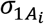 represent each participant’s observed variability on obstacle-present 2-target trials and obstacle-obstructed 1-target trials respectively, and 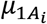 again represents each participant’s observed mean deflection, or safety margin, on obstacle-obstructed 1-target trials. Here *β* is a weighting parameter that controls the influence of the motor plans, or objectives, associated with optimization of movement timing and optimization of obstacle avoidance. Thus *β* = 1 indicates that participants prioritized movement timing for task performance and *β* = 0 indicates that participants prioritized obstacle avoidance for task performance. We assumed that 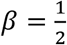 for the original PO model presented in Eq. 2 and for the corresponding predictions presented in Fig 5b,e, expressing the PO prediction as an equal balance of the two motor costs associated with the determinants of task performance.

We note that because the IMD that optimizes movement timing on 2-target trials (0°) would not be directly obstructed by the obstacle, it already affords some amount of safety margin around the obstacle. Thus the IMD that optimizes movement timing would also correspond to the IMD that optimizes obstacle avoidance if the expected safety margin during uncertainty is sufficiently small (i.e., *β* = 1). However, if the safety margin of the 0° movement is not large enough to optimize obstacle avoidance, then the IMD that optimizes both determinants of task success would indeed be based on a combination of the 0° IMD associated with optimization of movement timing and a unique, positively-valued IMD (i.e., skewed away from the obstacle) that optimizes obstacle avoidance based on variability. This feature of the PO model leads to corresponding predictions that evolve in a piece-wise linear fashion (as indicated by the horizontal and oblique line segments of the PO predictions shown in Fig 5b,e), but note that the safety margin estimate associated with optimization of obstacle avoidance went beyond the 0° direction for the vast majority of participants in both experiments (31/34).

Because the original MA and PO models assumed equal weighting of their respective model constituents, we assessed how their predictions might change if we allowed differential weighting between each model’s constituents. We thus fitted the MA and PO models above to the Expt 2a participant data (shown in Fig 5b) to estimate values for *α* and *β* that reflect participants’ own subjective valuations for the motor plans, or constituents, that comprise the MA and PO models. We then used the estimated parameters (*α* and *β*) to form refined predictions for the Expt 2b data.

We further note that the MA model predictions shown in Fig 5b,e were based on population-averaged estimates of *μ*_1*B*_, and analogously, the PO predictions shown in Fig 5b,e were based on population-averaged estimates of 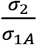. To appropriately assess the MA and PO models based on their participant-individualized predictions, we replotted the raw obstacle-present 2-target trial IMD data (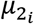; black dots in Fig 5b,e) such that the error between each datapoint 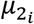 and the model predictions shown in Fig 5b,e reflects the error accrued from the *individualized* model predictions. We also note that the root-mean-square (RMS) of model prediction errors plotted in Fig 5a,d were determined from these individualized model predictions.

### Statistical Tests

T-tests were used for most statistical comparisons as indicated in the Results section. A one-sided t-test was used to compare obstacle-present versus baseline obstacle-free 2-target trial IMD data in Expt 2 (Fig 4b), and the remaining t-tests were two-sided. When we assessed nested forms of the full PO model in Expt 2 (see Eq. 3–4), we used F-tests. In all statistical tests, the significance level (alpha) was set to 0.01, and normality assumptions for the tests were verified with Kolmogorov–Smirnov tests.

## Acknowledgements

We thank Ashleigh Victoria Conroy-Zugel for help with experiments. This work was supported by a grant from the National Institute on Aging (R01 AG041878) to M.A.S.

## Data availability

All data acquired for this study used to analyze and generate all Figures have been deposited in OSF. DOI 10.17605/OSF.IO/S8A2W

